# High-resolution line-scan Brillouin microscopy for live-imaging of mechanical properties during embryo development

**DOI:** 10.1101/2022.04.25.489364

**Authors:** Carlo Bevilacqua, Juan Manuel Gomez, Ulla-Maj Fiuza, Chii Jou Chan, Ling Wang, Sebastian Hambura, Manuel Eguren, Jan Ellenberg, Alba Diz-Muñoz, Maria Leptin, Robert Prevedel

## Abstract

Brillouin microscopy (BM) can be used to assess the mechanical properties of biological samples in a 3D, all-optical, and hence non-contact fashion, but its weak signals require long imaging times and illumination dosages harmful to living organisms. Here, we present a line-scanning Brillouin microscope optimized for fast and high-resolution live-imaging of dynamic biological processes with low photo-toxicity. In combination with fluorescence light-sheet imaging, we demonstrate the capabilities of our microscope to visualize the mechanical properties of cells and tissues over space and time in living model organisms such as fruit flies, ascidians, and mouse embryos.

---

The fields of mechano- and physical-biology have burgeoned in the past decade, revealing that cell and tissue mechanics play an integral role in determining biological function^1^. While molecular components of cells can routinely be visualized with fluorescence microscopy in biology, assessing the mechanical, i.e. elastic and viscous, properties of living cells with similar spatio-temporal resolution is more challenging as current biophysical techniques^2–4^ used in the field exhibit intrinsic limitations. Recently, Brillouin spectroscopy has emerged as a nondestructive, label- and contact-free technique that can assess the visco-elastic properties of biological samples via photon-phonon scattering interactions^5–7^. By analyzing the frequency shift (*v_B_*) and linewidth (*Γ_B_*) of the light that is inelastically scattered from gigahertz-frequency longitudinal acoustic vibrations (phonons) in the sample, one can deduce the elastic and viscous properties of the sample, respectively. When coupled to a confocal microscope^8^, BM can achieve diffraction-limited resolution in 3D. Brillouin microscopy has enabled a wide range of applications in biology, including studies of intracellular biomechanics in living cells^9^, of ex-vivo^10,11^ and in-vivo tissues^12–14^ and their components^15^ (e.g., collagen, elastin, as well as biomaterials^16^) and early disease diagnosis^17^. Yet, despite its unique advantages, many 3D or live-imaging applications in biology remain practically out of reach, due to the fast dynamics or the photosensitivity of the biological samples involved. This is because virtually all BM implementations are based on single point-scanning (confocal) principle. This, together with the weak Brillouin scattering signal, typically leads to 2D imaging times in excess of tens of minutes to hours for large samples. At the same time, the confocal approach entails high and potentially phototoxic illumination dosages. While the improved signal levels of stimulated Brillouin scattering modalities^18,19^ have shown potential to decrease effective image acquisition time, the required high illumination dosages (~260mW in Ref.^18^) limit their use for live-imaging of sensitive samples over extended time-periods.

In this work, we present a Brillouin microscope specifically designed to address the limitation mentioned above and to enable a wider range of applications in cell and developmental biology. Our microscope is based on a line-scanning approach that enables multiplexed signal acquisition, allowing the simultaneous sensing of hundreds of points and their spectra in parallel. Furthermore, using a dual objective system at a 90-degree geometry together with near-IR illumination ensures minimal photodamage and phototoxicity. In contrast to previous proof-of-principle work^20^, our system design is specifically optimized for high spatio-temporal resolution as well as high signal-to-noise ratio (SNR), low phototoxicity and physiological sample mounting in order to enable long-term 4D mechanical imaging of highly sensitive biological specimen with sub-cellular resolution. An in-built, concurrent selective plane illumination microscopy (SPIM) fluorescence imaging modality allows for fluorescence guided Brillouin image analysis in 3D. We further implemented a GPU-accelerated, fast numerical fitting routine for real-time spectral data analysis and visualization, thus providing rapid feedback to the experimenter. We demonstrate the capabilities of our line-scan Brillouin microscope (termed LSBM) by live-imaging the 3D mechanical properties of developing *Drosophila melanogaster*, *Phallusia mammillata* and mouse embryos over an up to ~180×165×170μm field-of-view (FOV) with down to 1.5μm spatial and down to 2min temporal resolution.

The design of LSBM imaging system is conceptually shown in **Fig. 1** and is based on an inverted SPIM configuration^21^ with high (0.8) numerical aperture, as well as a narrowband (50kHz) yet tunable 780nm diode laser (**Online Methods** and **SI Fig. 1**). This wavelength significantly reduces phototoxic effects compared to 532nm laser lines that are more commonly used in confocal BM^8,14,22^. Moreover, the near-IR wavelength does not interfere with the excitation spectra of common fluorescence labels. The tuneability of the laser allows to stabilize the frequency by locking to well-defined atomic transitions by means of absorption spectroscopy (D2 line of Rb^87^) and to employ a gas cell as an ultra-narrowband notch filter^13,23^ for the suppression of inelastically scattered Rayleigh light to within ~80dB. Such high background suppression is critical for imaging biological tissues. To ensure the necessary high spectral purity of the diode laser, we built a custom narrowband filter consisting of a Bragg grating and a cavity-stabilized Fabry-Perot interferometer to suppress amplified spontaneous emission (ASE) noise below ~90dB (**SI Fig. 2**, **Online Methods**).

**Figure 1.**
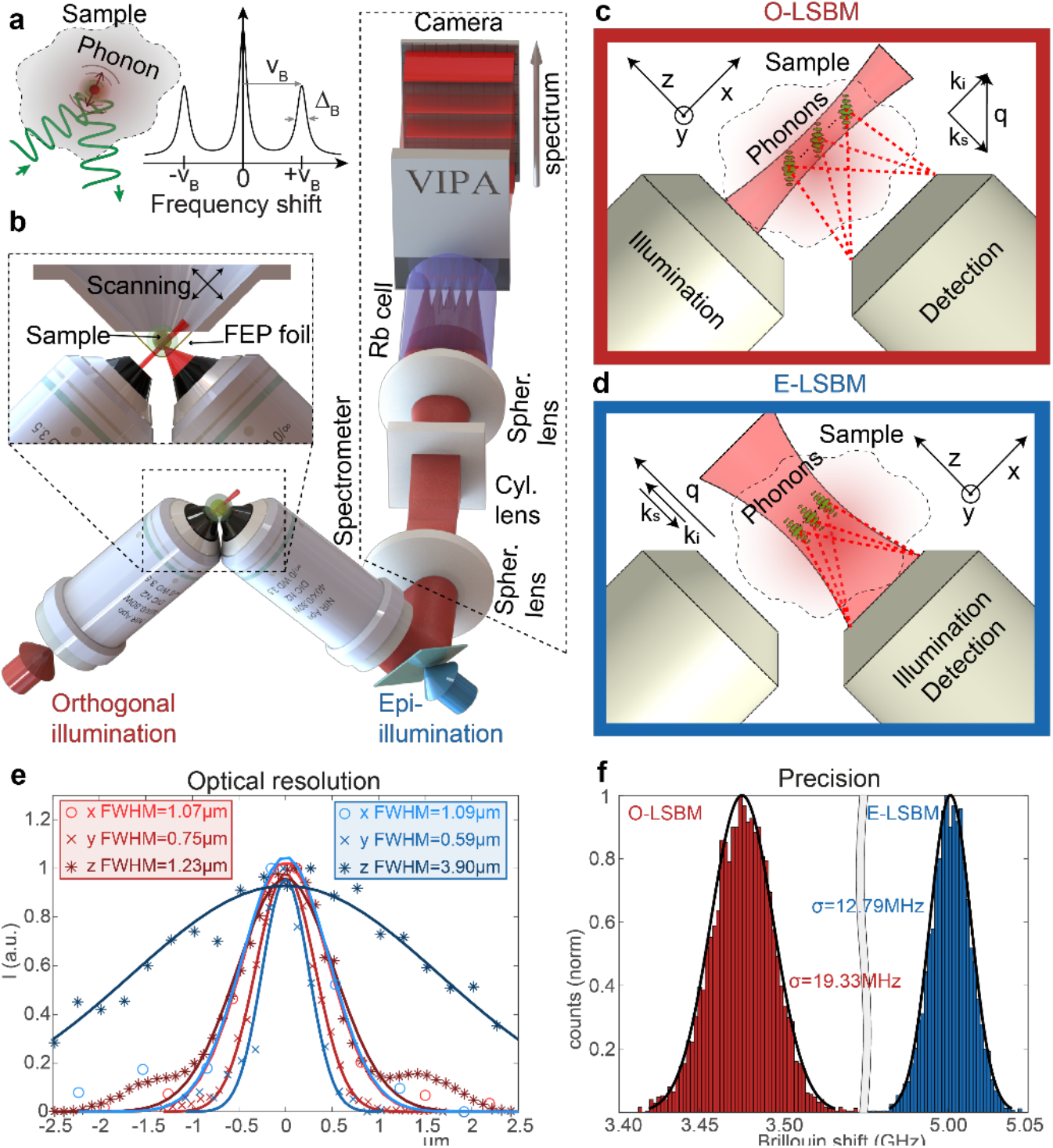
Schematic and characterization of the Brillouin line-scanning microscope. **(a)** Principle of Brillouin scattering: a small portion of the incident laser light is scattered by spontaneous thermal-induced sound waves (phonons) and gives raise to the Brillouin spectrum (**b**). LSBM setup: The light is delivered/collected by two objectives, mounted in an inverted 90 degrees configuration. The sample is placed in a chamber (attached to a 3-axis translational stage for scanning) which is separated from the surrounding immersion liquid by a thin FEP foil (inset). The light collected from the sample is redirected to the spectrometer (dashed box) which consists of a combination of spherical and cylindrical lenses to generate the correct beam profile to be input on the Virtually Imaged Phased Array (VIPA) that disperses the light on the camera. A pure Rb^87^ cell rejects the elastically scattered (Rayleigh) light. Scattering geometry for the **(c)** orthogonal and **(d)** epi configurations of the LSBM: in the orthogonal geometry the illumination direction (**k_i_**) is perpendicular to the detection (**k_s_**), consequently the phonon direction (**q**) is at 45deg compared to both of them. In the epi geometry only a single objective is employed and the illumination direction is parallel to both the detection and the phonon direction. The convention for the x, y, z axes is the same used in all the figures. **(e,f)** Characterization of the LSBM in terms of **(e)** spatial resolution (measured by Rayleigh-scattered light from 0.5 μm beads) and **(f)** spectral precision (determined by calculating the standard deviation of multiple measurement of Brillouin shift in water in typical experimental conditions) for both the orthogonal and the epi geometries (color-coded in red and blue, respectively). For details concerning the SPIM modality integration, see Supplementary Fig. 1.

The opto-mechanical design of the microscope allows operation in two alternative geometries for Brillouin imaging (**Fig. 1c,d** and **SI Fig. 1**): **(1)** Either the illumination and detection axis can be separated in a 90-degree fashion (termed ‘orthogonal line’ or O-LSBM, **Fig. 1c**), similar to SPIM, in order to minimize total illumination dosage for volumetric imaging, or **(2)** the sample can be illuminated by a focused line in a 180-degree backscattered fashion (termed ‘epi-line’ or E-LSBM) to mitigate effects from scattering and optical aberrations (**Fig. 1d**). In both cases, the optical parameters of the microscope and the overall performance of the spectrometer were optimized for high spatial resolution (matched to typical phonon lengths in biological material^24^), background suppression as well as the comparatively large FOV (~200μm) required for common model organisms in biology (**Fig. 1e, SI Fig. 3-5**). To maintain homogenous focusing of the illumination line across a ~200μm FOV in the O-LSBM configuration, we employ an electrically tunable lens (ETL) (**SI Fig. 6** and **Online Methods**).

The microscope’s sample stage incorporates a Fluorinated Ethylene Propylene (FEP) foil to physically isolate the specimen chamber from the objective’s immersion media which allows the use of microdrop cultures required for longitudinal embryo imaging (**Fig. 1b**). Furthermore, we designed a custom miniaturized incubation chamber for full environmental (temperature, CO_2_, O_2_) control (**SI Fig. 1c-d**). To analyze the multiplexed (100+) spectra of the LSBM in real-time and ensure high precision spectral measurements, we implemented a GPU-enhanced numerical fitting routine tailored to the lineshape of our experimental spectra^25^, yielding a >1000-fold enhancement in processing time and thus real-time spectral data analysis (**SI Fig. 7, Online Methods**). Overall, the line scanning Brillouin imaging system achieves SNR (~10dB) and spectral precision (<20MHz) that are comparable to single-point scanning BM implementations^9,14,26^, and should thus be suitable for imaging of biological samples at much reduced illumination dose (**Fig. 1f, SI Fig. 4d**).

To demonstrate the capabilities of the LSBM we captured longitudinal and 3D mechanical information of various developing organisms on physiologically relevant timescales. First, to highlight the capacity of the LSBM to detect mechanical changes of highly dynamic processes such as organismal-level morphogenesis, we recorded two fast and widely studied events in *Drosophila* gastrulation, ventral furrow formation (VFF) and posterior midgut invagination (PMG) (**Fig. 2**). VFF and PMG are fast tissue folding events progressing in the minutes timescale, both driven by actomyosin but with differential geometry of their contractile domain (rectangular in VFF, circular in PMG)^27^ (**Fig. 2a-b,e-f**). During early VFF, cells with mesodermal fate (marked by the expression of the transcription factor Snail) located on the ventral side of the embryo undergo complex changes in cell geometry that will enable the complete invagination of these cells in ~15min^28^ (**Fig. 2a,b**). By using the LSBM in epi-line geometry (E-LSBM) we were able to record both fluorescence SPIM as well as Brillouin shift maps of VFF over a FOV of ~22×170×71μm (z-increment of 1.5μm) and of PMG over an FOV of ~83×141×46μm (z-increment of 2.5μm), both sampled with a volume time-resolution of ~2min (**Fig. 2c,f** and **SI Video 1-2**). This equals ~10sec per 2D image slice or an effective pixel time of ~1ms inside live biological tissues, which represents an approximate 100- (20-)fold improvement compared to previous spontaneous (stimulated) Brillouin scattering microscopes, respectively, at >10-fold lower illumination energy per pixel^7^. Analysing the volumetric Brillouin data based on fluorescence membrane labels, we found that the average Brillouin shift of cells engaged in VFF transiently increases within the presumably mesoderm by 10-20MHz from the point of apical constriction initiation (0’) until mesoderm invagination is completed (p=0.039, **Fig. 2c,d, SI Video 1**). We further imaged PMG invagination, in which the cells form a circular contractile domain (**Fig. 2e,f, SI Video 2**). Similar to the observations during VFF, the average Brillouin shift within cells engaged in tissue folding also increased during PM (**Fig. 2f**). This suggests that during tissue folding a higher Brillouin shift is a common feature independent of the geometry of the contractile domain. No photodamage or - toxicity was observed at <~20mW of average laser power, and viability assays showed that all embryos (n=3) imaged progressed to the first larval stage (24hpf).

**Figure 2.**
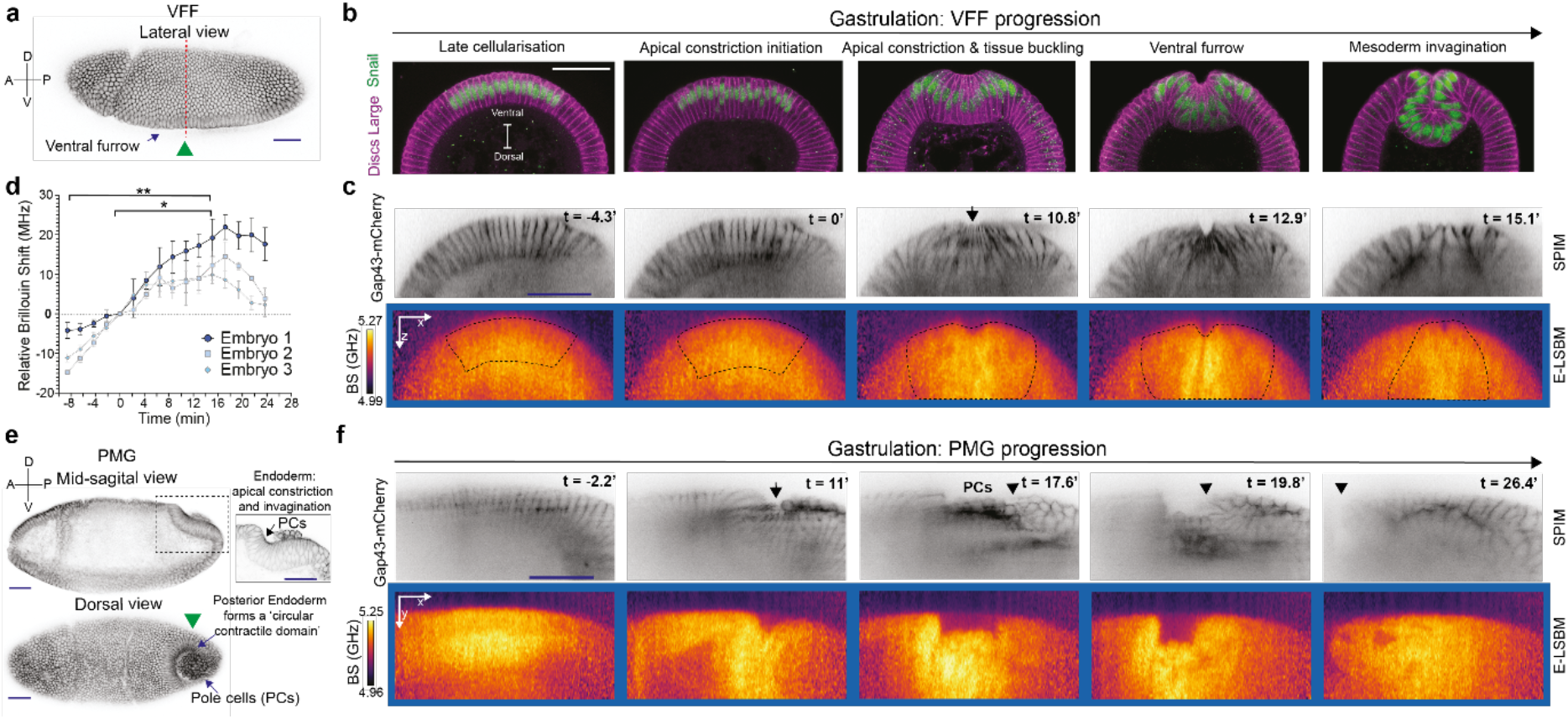
Cells undergoing tissue folding events during *Drosophila* gastrulation display high Brillouin shifts. **(a)** Superficial view of a fixed *Drosophila* embryo undergoing ventral furrow formation (VFF, arrow). The embryo expresses the membrane marker Gap43-mCherry to visualise cell outlines. Green arrowhead indicates the direction of Brillouin illumination which here is orthogonal to the apical surface of cells. Red segmented line indicates the approximate location of physical cross-sections shown in **b** and **c**. **(b)** Stages of VFF progression shown in physical cross-sections of fixed embryos, stained with a Snail antibody (green) to label the nuclei of cells engaged in VFF and a Discs Large antibody (magenta) to label cell outlines. (**c**) Top: Images from a SPIM recording of an embryo expressing Gap43-mCherryto mark cell membranes; bottom: median-projected Brillouin shift maps from the same embryo at the same timepoints and positions as shown above (representative embryo of n = 3 total). Dotted line encloses the presumable mesoderm region within the Brillouin shift maps used for quantifications in **d**. The apical side of cells faces the outer surface of the embryo. **(d)** Quantification of Brillouin shift averaged over 18 cells engaged in apical constriction and invagination during VFF progression, as inferred from SPIM slices (dotted line in **c**). Changes in Brillouin shift are shown relative to the Brillouin shift at time-point 0’, which corresponds to the initiation of apical constriction. Data points are average Brillouin shift within contractile domain, error bars denote S.D. over different slices. Statistical significance was evaluated using a paired-one-way ANOVA test (F = 44.29, p = 0.066), followed by a post-hoc multiple comparison test (FDR corrected, α = 0.05, q-value = 0.1); * p = 0.039, ** p = 0.0054. **(e)** Posterior midgut invagination (PMG) in embryos at a similar stage as in **a**. (Top) mid-sagital section; inset: higher magnification of the posterior midgut region showing apical constriction and invagination during PMG formation; (bottom) superficial-dorsal view a *Drosophila* embryo showing the formation of the circular contractile domain of the PMG that encloses the Pole Cells (PCs, arrow). Green arrowhead indicates the direction of Brillouin illumination which here is orthogonal to the lateral surface of cells. **(f)** Representative SPIM images (top) and median-projected Brillouin shift maps (bottom) at five timepoints during invagination (out of n = 3 total). Arrowhead indicates the displacement of the posterior end during PMG invagination. The initiation of apical constriction was set as 0’ time point. Images in Gap43-mCherry and Brillouin panels are median projections of 3 slices of the re-sliced ROI (see Methods). Timescale is minutes. Scale bars are 50 μm.

In order to further explore the potential of the LSBM to collect high-resolution 3D mechanical data from developing organisms, we imaged the ascidian *Phallusia mammillata*, a simple chordate with highly conserved stereotypic cell lineages and cellular organisation (**Fig. 3)**. Here, we used the orthogonal line configuration of the microscope (O-LSBM) to collect high-resolution, sub-cellular elasticity information in the early embryo, over a FOV of 165×186×172μm (z-increment of 2.5μm) and within ~17min, shorter than the average time in between developmental stages of this animal at 18°C (**Fig. 3b,c**). We observed a perinuclearly localized, high Brillouin signal within the B5.2 cells in the late 16-cell stage (**Fig. 3e,f**). This subcellular region is known to have a dense microtubule bundle structure driven by the centrosome attracting body (CAB)^29^ (**Fig. 3d**). We also applied the epi-line configuration (E-LSBM) to a later developmental stage (late tailbud I), in order to study tissue level differences of mechanical properties in 3D (**Fig. 3g-j**). Here, with fluorescent SPIM imaging of the lipophilic dye FM4-64 as a membrane marker, we were able to distinguish between the epidermis, central nervous system (CNS) and mesoderm/endoderm cells and observed differences in Brillouin shift between them (**Fig. 3j**), which possibly suggest that the different nature and organization between the epithelial (epidermis and CNS) and mesenchymal cell types (endoderm and mesoderm) results in differential tissue mechanics. These experiments also demonstrate that LSBM imaging can be used over long periods (here 14 hours; **SI Video 3-4**), thus enabling tracking changes in mechanical properties driven by cellular differentiation over time.

**Figure 3.**
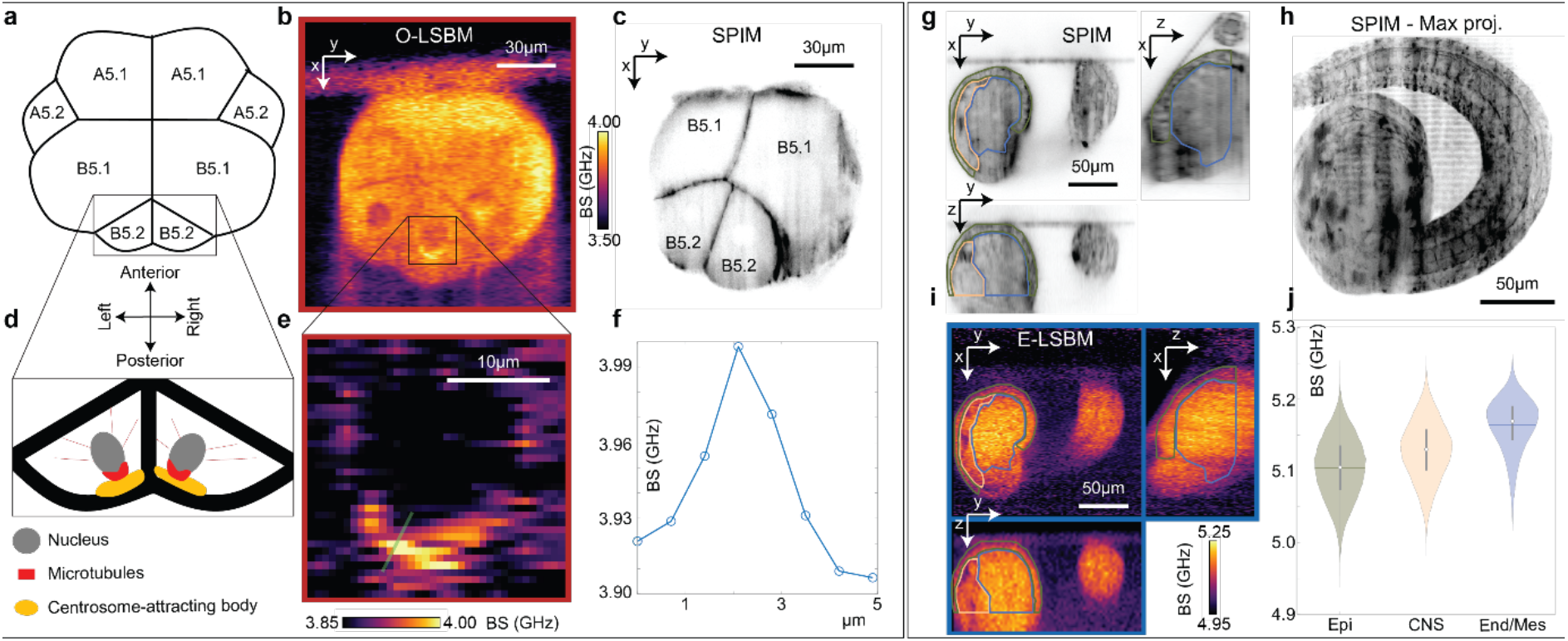
LSBM imaging of *Phallusia mammillata* shows mechanical differences on subcellular scale and between tissue types over large volumes in 3D. **(a)** Schematic of the vegetal hemisphere of a late 16 cells stage *Phallusia* embryo, showing the bilateral symmetry. **(b)** Exemplary O-LSBM image and **(c)** corresponding SPIM image (membranes labelled with lipophilic dye FM4-64) of the same stage as shown in **a** (out of n = 2 total). **(d)** Cartoon of the machinery for asymmetric division in two B5.2 germline progenitor cells: a thick microtubules bundle directed to the centrosome-attracting body forms on the posterior side of the nucleus. **(e)** Zoom-in of the dark gray square in panel **b** showing a high Brillouin shift in the vicinity of the nucleus that is indicative of a microtubule bundle. **(f)** Line profile along the line shown in panel **e**. **(g-j)** Imaging of a *Phallusia* embryo at the tailbud stage (out of n = 3 total). **(g)** Orthogonal views and **(h)** maximum intensity projection (MIP) from the SPIM volume (195μm x 191μm x 111μm) of membranes labelled with lipophilic dye FM4-64 of a *Phallusia* embryo at the tailbud stage (late tailbud I) **(i)** Corresponding orthogonal views, acquired with E-LSBM modality, of the volume in panel **g**. **(j)** Violin plot showing the Brillouin shift differences of three distinct tissue regions (Epidermis, Central Nervous System (CNS) and Endoderm/Mesoderm) manually-segmented in 3D according to SPIM data in **g**. The green, yellow and light-blue outlines in panels **g** and **h** show the segmented regions for Epidermis, CNS and Endoderm/Mesoderm respectively. BS, Brillouin shift.

Finally, to test the low-photo-toxicity of the LSBM approach in a further organism, we imaged the developing mouse embryo, from the 8-cell stage (E2.75) to the late-blastocyst (E4.5), covering a 46-hour time-span (**SI Fig. 8**). We acquired 3D Brillouin time-lapse volumes over a ~100×125×110μm FOV (z-increment of 1.5μm) within ~11-17min and at 75-90min timeintervals as well as simultaneous SPIM data which allowed cell tracking via fluorescently labelled nucleus (H_2_B-mCherry) and membrane (Myr-Palm-iRFP). Despite the embryos’ notorious photo-sensitivity, no photodamage or -toxicity was observed at <~20mW of average laser power as confirmed by the morphology, dynamics, cell number and cell fate of the imaged embryos resembling those of control embryos (**SI Fig. 8c,d**). This represents a significant improvement over standard confocal BM employing 532nm illumination, where embryo death was observed following a few 2D image acquisitions (**SI Fig. 9**).

To summarize, here we presented a multi-modal line-scan Brillouin microscope for fast and high-resolution 3D imaging of mechanical properties of highly sensitive living organisms in biology. In particular, we have shown for the first time that entire multicellular organisms can be successfully imaged with Brillouin microscopy at high spatial resolution in 3D and over extended time-periods without apparent photodamage. Compared to alternative Brillouin scattering approaches and implementations^7,9,13,14,18^ this represents a >20-fold improvement in terms of imaging speed, at >10-fold lower illumination energy per pixel without sacrifices in measurement precision^7^.

These advances have allowed us to demonstrate that LSBM can detect changes in mechanical properties at the cell and tissue scale that occur for example during early development in a broad range of organisms. Specifically, we reveal that a higher Brillouin shift is a common feature of tissue folding, independent of the geometry of the contractile domain (**Fig. 2c,f**). We were also able to visualize subcellular structures with different mechanical properties that likely relate to the cytoskeleton (**Fig. 3b-e)**, as well as differential tissue mechanics deep inside an embryo (**Fig. 3g-j)**, without the need for tissue sectioning (and its concomitant artefacts) or the invasive injection of micro-particles or -droplets.

The fact that our LSBM system can easily switch between two scattering geometries allows choosing the best performance for a given application. While the O-LSBM provides the lowest photo-damage burden and best axial resolution, optical sample accessibility from two sides must be ensured, and optical refraction effects between the sample and medium can affect image quality. The latter could in principle be mitigated by refractive index matching or adaptive optics methods^30^. In contrast, the E-LSBM geometry is inherently insensitive to these effects, and is thus better suited for more scattering or heterogeneous samples, yet has slightly lower axial resolution and a higher illumination light burden for large volumes. In the future, the symmetric design of the microscope described here can be employed for rapid, dual-view imaging by optically switching the illumination and detection paths, an approach that could mitigate shadowing effects and improve resolution through data fusion^31^.

To conclude, we expect that the low phototoxicity and significantly improved acquisition speed of LSBM will open up exciting new possibilities for the field of mechanobiology. In particular, we anticipate enticing applications in imaging mechanical changes during development together with e.g. cytoskeletal or cell-fate specific fluorescence reporters, or by correlating 3D mechanical to molecular, genetic or ultra-structural datasets^32^. This would likely shed new light on the complex interplay between genetics, biochemical signaling and the role of biomechanics in animal development.

## ACKNOWLEDGMENTS

We would like to thank the mechanical and electronic workshop at EMBL Heidelberg for help; Takashi Hiiragi and Lars Hufnagel for support in early stages of the project; Isabell Schneider and Koki Watanabe for help with mouse imaging experiments; Sergio Lembo for help with confocal imaging of the stained embryos; EMBL LAR for mouse husbandry and ascidian support. We thank Fiona Paul Ukken for the generation of the *Drosophila* Snail antibody and Elisabeth Vogelsang for producing physical cross-sections of *Drosophila* embryos. We further thank Stefano De Renzis (EMBL) and Natalia Bulgakova (University of Sheffield) for providing the sqh-Gap43::mCherry and EB1-GFP transgenic line, respectively. A.D.-M. was supported by the Deutsche Forschungsgemeinschaft (DFG) research grants DI 2205/2-1 and DI 2205/3-1. The lab of M.L. was funded by EMBO and DFG grant LE 546/12-1. R.P. acknowledges support of an ERC Consolidator Grant (no. 864027, Brillouin4Life), and the German Center for Lung Research (DZL). R.P. and A. D.-M. acknowledge funding from the COST Action CA16124 (‘BioBrillouin’). This work was supported by the European Molecular Biology Laboratory.

## AUTHOR CONTRIBUTIONS

R.P. conceived the project and designed the imaging system together with C.B. C.B. built the imaging system, and wrote control software. C.B. performed experiments and analyzed data together with C.J.C, J.M.G., and U.F., under guidance of A.D.-M., M.L and R.P.. L.W. designed mechanical components. S.H. wrote the GPU spectral analysis code. J.M.G.. provided transgenic fly lines and analyzed data, under guidance of M.L.. M.E. provided and injected transgenic mouse embryos under guidance of J.E. R.P. led the project and wrote the paper with input from all authors.

## Methods

### Line-scan Brillouin imaging setup

A detailed schematic of our LSBM is shown in Supplementary Fig. 1. The laser is an amplified tunable diode laser (TA pro, Toptica) locked to the D2, F=2 absorption line of Rb^87^ (780.24nm). To further clean up the laser’s spectrum from ASE noise, we designed a custom-built narrowband double-pass optical filter^33^, consisting of a Polarising Beam Splitter (PBS) followed by a tunable Fabry-Perot cavity (FSR 15GHz, Finesse ~60, LightMachinery), an holographic diffraction grating (NoiseBlock, Coherent), a quarter waveplate and a mirror. A beam sampler (BSF10-B, Thorlabs) sends ~5% of the light to a photodiode whose signal is fed into an Arduino; a self-written code maximizes the transmitted intensity, to correct for longtime drift of the cavity (Supplementary Fig. 2a). The filter enables a total suppression of the integrated ASE noise down to ~-90dB, more than 40dB below the original ASE level of the laser (Supplementary Fig. 2c,d). After the filter, the light is coupled into a polarization maintaining (PM) fiber and delivered to the main microscope. Here, a collimator (LPC-OT-780-5/125-P-3-16AC-40-3A, OZ Optics) produces a collimated beam with a diameter of 2.77mm (1/e^2^, theoretical), that corresponds to a focused beam of 1.79μm (1/e^2^, theoretical) after the objective. The beam then passes through an electrically tunable lens (ETL) (EL-10-30, Optotune) coupled with a negative offset lens (LC1258-B, Thorlabs). This allows axial scanning of the focus position after the objective (Supplementary Fig. 6). After the tunable lens, a flip mirror allows to switch between the epi-line and orthogonal-line illumination: the separate optical path for the epi-line illumination consists of a 40mm lens (LA1422-B, Thorlabs), that collimates the light out of a PM fiber (P3-780PM-FC-2, Thorlabs), providing a beam of diameter 8.7mm (1/e^2^, theoretical) (filling the back focal aperture of the objective, which is 8mm); a 200mm cylindrical lens (LJ1653RM-B, Thorlabs) generates the focused line in the focal plane of the objective with an extension of ~185μm (1/e^2^) (Supplementary Fig. 6). A half waveplate (WPH05M-780, Thorlabs) is used to rotate the linear polarization of the illumination light to match the required direction for the two geometries. Two mirrors, conjugated to the back and the front focal plane of the objective (by means of two 200m lenses: #33-362 and #49-364, Edmund), respectively, allow for fine position and angle adjustment of the beam inside the sample in order to aid in the alignment between the illumination and detection objectives.

For the fluorescence SPIM modality, we utilize a laser box providing several laser lines for fluorescence excitation (405nm, 488nm, 560nm, 640nm). The fluorescence excitation is introduced into the setup by a single mode fiber (P1-460Y-FC-1, Thorlabs) whose output is collimated by an achromatic lens (f=4mm), producing a ~0.8mm (1/e^2^, theoretical) beam. A fiber polarisation controller (FPC030, Thorlabs) is used to adjust the polarisation and a polariser rejects potential residual non-linear polarisation. The excitation light is then coupled into the same optical path as the Brillouin illumination by means of a dichroic mirror (FF765-Di01, Semrock). Similarly, to the Brillouin illumination path, two lenses (#67-159, Edmund and #49-364, Edmund, shared with the Brillouin path) and mirrors (not shown in Supplementary Fig. 1) allow for beam adjustment. A 18mm cylindrical lens (#68-041, Edmund) is used to generate the light sheet. The corresponding thickness and lateral extent of the light sheet inside the sample are ~4μm and ~200μm (1/e^2^, theoretical), respectively.

The detection and illumination objectives are identical (Nikon, 40x, 0.8NA, MRD07420, water immersion) and mounted in an inverted V configuration to facilitate imaging of small (~200μm) samples in physiological conditions^21^ (Fig. 1b and Supplementary Fig. 1). The sample is placed in a dedicated, custom chamber that utilizes a 12.7μm-thick FEP foil (RD-FEP050A-610, Lohmann) to separate the immersion water of the objectives from the sample. The sample’s environment (medium) is temperature controlled via a custom-built heating element and PID controller, and CO_2_ is added via a custom-built gas-mixer (Supplementary Fig. 1c,d). For 3D image acquisition, the chamber containing the sample is scanned by three single-axis translational stages (SLC-2430, SmarAct).

For the epi-line Brillouin geometry (E-LSBM), a polarising beam splitter (PBS) (BPB-12.7SF2-R400-800, Lambda), mounted on a magnetic base for easily switching between the two modalities, followed by quarter waveplate (WPRM-25.4-12.7CQ-M-4-780, Lambda) reflects the light towards one of the two objectives; the Brillouin signal is collected by the same objective and transmitted by the PBS towards the spectrometer (the polarisation is rotated by 90 degrees due to double pass at the quarter waveplate). A polariser is added before the other objective (used for SPIM excitation) which is required in order to block the backwards-scattered un-polarised fluorescence light that would otherwise be reflected by the PBS and generate a bright line at the center on the FOV of the SPIM camera.

The fluorescence and Brillouin signals are split via a dichroic mirror (FF765-Di01, Semrock), reflecting the fluorescence emission. A tube lens (MXA20696, Nikon) focuses the fluorescence image on the SPIM camera (Zyla sCMOS, Andor); a motorized filter wheel (96A361, Ludl) allows for the selection of the proper emission filter. The Brillouin signal is instead transmitted by the dichroic and sent to the spectrometer (Supplementary Fig. 3). Here, first two spherical lenses (LA1256-B and LA1979-B-ML, Thorlabs) relay the BFA of the detection objective and a slit (VA100/M, Thorlabs), placed in the focal plane of the first lens, ensures confocality in the direction perpendicular to the illumination line. After the relay, inelastically scattered (Rayleigh) light is filtered by a 150mm-long gas cell filled with pure Rb^87^ (SC-RB87-(25×150-Q)-AR, Photonics Technologies). The cell is wrapped into heating foil (HT10K, Thorlabs) and a self-built electronic board (based on an Arduino Mega) controls the temperature of the cell via PID. After the Rb gas cell, a combination of an achromatic lens (#49-794, Edmund), a cylindrical lens (#36-231, Edmund) and a spherical lens (#69-513, Edmund) relay and focus the light on the entrance slit of the VIPA (OP-6721-6743-9, LightMachinery). In-between the second spherical lens and the VIPA there is an additional 75mm-long Rb^87^ cell (SC-RB87-(25×75-Q)-AR, Photonics Technologies).

After the VIPA two cylindrical lenses (LJ1267L1-B and LJ1629L2-B, Thorlabs) focus the light on the EM-CCD camera (iXon DU-897U-CS0-BV, Andor), preceded by a bandpass filter (FBH780-10, Thorlabs) to block ambient light. These lenses are chosen to ensure proper spatial and spectral sampling (0.7μm/px and ~0.25 to ~0.5 GHz/px). The optical layout and performance are shown in Supplementary Figs. 3-5. To synchronize the ETL with the EMCCD acquisition, the fire output of the EMCCD (high when the camera is exposing) is used to trigger a DAQ (PCI-6221, National Instruments) that generates the voltage ramp driving the ETL. The entire data acquisition is controlled by a self-written LabView program and the image reconstruction is performed with a Matlab script, that provides an interface to the GPU-fitting library (see section Brillouin line scan spectral analysis).

### LSBM system characterization and image acquisition

To quantify the optical resolution of the Brillouin spectrometer, we embedded 0.5μm fluorescent beads (TetraSpeck, Thermo Fischer) in 1% agarose and recorded the intensity of Rayleigh scattering while scanning the sample (Fig. 1e, Supplementary Fig. 4a,b). To quantify the spatial pixel size we moved the bead along x and performed a linear fit (SI Fig 4c). To measure the spetrometer’s spectral resolution, we collected the Brillouin signal from scattering intralipid solution, remapped and summed the spectra (see section Brillouin line scan spectral analysis) and performed a Lorentzian fit on the Rayleigh signal. The ASE was measured with the setup shown in Supplementary Fig. 2b that consists of the addition of a diffraction grating to the Brillouin spectrometer to have an extended FSR while retaining high spectral resolution. The Rb cells suppress the laser light, thus allowing for a very high dynamic range (>90dB) needed for these measurements.

### Brillouin line scan spectral analysis

Each spectral line (corresponding to the light scattered from a point along the illumination line) is analyzed separately. The processing of this spectral data is then done in 3 main steps: First, the spectrum is remapped into a linear frequency space. Then, the signal from at least three different orders is summed in order to increase the effective SNR before the Stokes and anti-Stokes peaks are fitted separately. To speed up the entire processing via parallelization, we decided to implement this analysis pipeline with custom GPU code written in CUDA. The linear remapping (pixel-to-frequency conversion) depends only on the position of the Rayleigh peak and the FSR of the VIPA (15.15GHz). Here, the FSR was measured from the calibration of the spectrometer with Rb D2 hyperfine lines of known frequency (Supplementary Fig. 5). For the remapping we use spline interpolation whenever the target frequency sampling does not match the frequencies of individual pixels. Each line of the image can be processed independently, allowing for this operation to be executed in parallel on the GPU. After this preprocessing step, the individual spectra are fitted over a user selected range using GPUfit^34^. Lorentzian, quadratic and broadened Brillouin lineshape^25^ functions were custom added into this library. In addition to the main Brillouin peak position and linewidth, also other fitting parameters, such as amplitude, SNR, goodness of fit, and number of iterations can be retrieved. The reported Brillouin shift is the average between the frequency shift of the Stokes and anti-Stokes peak.

In order to benchmark our GPU pipeline, we created a synthetic Brillouin spectral signal, which we then processed with either the standard, CPU-based Matlab-script, or our custom GPU code. On our PC (Intel Core i9-11900K, 64GB RAM, and Nvidia GeForce GTX 1050 Ti), the GPU processing took about 10ms regardless of the dimension of the image, while the purely Matlab based pipeline took more than ~1sec, and scaled linearly with the image size (Supplementary Fig. 7a). Both processing methods give similar results, with negligible (~1MHz) difference between the fitted peak frequencies and the real value (Supplementary Fig. 7b).

### *Drosophila* embryo imaging

#### Live imaging

*Drosophila* live imaging was performed with embryos expressing Gap43::mCherry to label cell membranes and EB1-GFP to mark microtubules plus ends^35^. Embryos for Brillouin imaging were prepared as follows: 1-hour synchronised egg collections were performed at 25°C and embryos were allowed to develop a further hour at 25°C. Next, Halocarbon oil (27S, Sigma) was added on the agar plate carrying the collected embryos to enable accurate embryonic staging through visual inspection of morphological features under a binocular (Zeiss). Embryos in Stage 5a-b^36^ were hand-selected, dechorionated in Sodium Hypochlorite (Standard Bleach, 50% in H_2_O) for 90 seconds, and washed using PBS 1X. Finally, embryos were glued (with Heptane-Glue) to the outer surface of the FEP film in the desired orientations to enable the collection of data from different angles. Image acquisition was initiated 5 minutes after the cellularising front of the blastoderm cells passed the basal side of the nuclei. The temperature of the imaging chamber was set to 22°C.

#### Drosophila Snail antibody generation

A construct encoding an N-terminal gluthathione S-transferase (GST)-tagged portion of the *Drosophila* Snail protein (encompassing aminoacids 1-200 of Uniprot ID P08044) was expressed in *E. coli* and purified from lysates of bacteria expressing the Snail-GST fusion protein using beads with immobilised glutathione. The fusion protein was eluted from beads using reduced glutathione and used for immunising a rabbit. The immunisation and collection of pre-bleeds and test bleeds was performed by Cocalico Biologicals Inc. The antiserum was tested for specificity and titer by immunostainings, immunoprecipitation and Western blot analysis.

#### Immunohistochemistry (IHC)

*Drosophila* whole mount embryos were fixed and stained as previously described^37^. In Fig. 2a and 2e, Gap43-mCherry transgenic embryos were stained using an mCherry antiserum (rabbit, 1:1000, Abcam ab167453). In Fig. 2b, *w^1118^* embryos were co-stained with the anti-Snail antibody described above (Rabbit, 1:500) and an anti-Discs Large antibody to label the lateral membrane of cells (mouse, 1:50, 4F3, DSHB). Physical cross-sections across half of embryonic length were produced on these stained embryos as previously described^38^.

#### Image processing and automated image analysis

To allow for SPIM fluorescence image guided analysis, the Brillouin time-lapse images were first re-scaled (bilinear interpolation) in ‘x’ and ‘y’ using a 2.5138 factor to match them to the dimensions of the fluorescence images. The fluorescence time-lapse was manually aligned with the Brillouin time-lapse using the y-coordinate half of the diameter of the fluorescence image (450 pixels) and using the colocalisation of the furrow as a landmark in ‘x-coordinate’ (either the VFF or PMG Furrow). The aligned fluorescent time-lapse was cropped using the Brillouin dimensions to stack the two time-lapses and produce an overlay image. For VFF image processing and quantification, for each analysed embryo the overlaid time-lapse was re-sliced and 3 median-projections of two slices each were used as embryonic replicates. To conduct the automated image analysis, we manually generated masks that included an 18-cell domain presumably containing the mesoderm (Fig. 2b, ‘Late cellularisation’, Snail-positive cells). The region enclosed by the mask was determined by taking 9 cells in each direction (left and right) from the center of the furrow. These masks were loaded by a Matlab script that filtered the Brillouin shift maps within each corresponding time-point for each replicate and for each embryo. The average Brillouin shift within the filtered, presumably mesodermal, domain and the corresponding standard deviation were calculated per embryo and plotted against time, taking time = ‘0‘ as the first sign of apical constriction. Image quantification was performed on 3 independent embryos undergoing VFF.

#### Statistical Analyses

The statistical significance of the differences of the average Brillouin shift within the filtered domain across the 3 independent embryos at timepoints −8.6’, 0’ and 15.1’ was analysed by applying a paired one-way ANOVA, which was followed by a post-hoc Multiple Comparison Test (FRD correction, Methodology: Two-stage-step-up method of Benjamini, Krieger and Yekutieli, q-value = 0.1). Sphericity was not assumed (equal variability of differences). Statistical significance was set at a = 0.05. Normality of the distribution of averages at each evaluated time point was analyzed using a Shapiro-Wilk test. Statistical analyses were performed using Prism GraphPad V8.

### *Phallusia* imaging

#### Live Imaging

Adult *Phallusia mammillata* were provided by the Roscoff Marine Biological Station (France) and kept at 17°C under constant illumination. *In vitro* fertilised embryos were prepared as described in Ref.^39^. Individual membranes were labelled with and imaged in artificial sea water with 5μg/ml lipophilic dye FM4-64 (Invitrogen). The imaging chamber of our LSBM system was kept at 12C and 17C, respectively in Fig. 3b and Fig. 3i.

#### Image data analysis

The registration of the SPIM and Brillouin images for analysis was done similarly to the *Drosophila* embryo, i.e. the Brillouin image was upscaled (bilinear interpolation) to match the pixel size of the fluorescence image. A rectangular area of the same size as the Brillouin image was manually aligned around the center of the fluorescence image (using the edges of the embryo, visible in both modalities, as a guide) to match the Brillouin image. The boundaries of different tissue types were manually drawn, slice by slice, by looking at the morphology of the cells in the fluorescence image. The segmentation masks were then imported in Matlab and used to generate the violin plot in Fig. 3j.

### Mouse embryo imaging

#### Recovery and culture

Animal work was performed in the animal facility at the European Molecular Biology Laboratory, with permission from the institutional veterinarian overseeing the operation (ARC number 2020-01-06RP). The animal facilities are operated according to international animal welfare rules (Federation for Laboratory Animal Science Associations guidelines and recommendations). (C57BL/6xC3H) F1 mice from eight-weeks of age onwards were used. Embryos were recovered from superovulated female mice mated with male mice. Superovulation was induced by intraperitoneal injection of 5 international units (IU) of pregnant mare’s serum gonadotropin (Intervet Intergonan), followed by intraperitoneal injection of 5 IU human chorionic gonadotropin (Intervet Ovogest 1500) 44–48 h later.

#### Imaging

Sample holder was covered with pre-equilibrated (5 % CO2, 37°C) G-1 Plus medium (~30 μl) and covered with pre-equilibrated Ovoil (~150 μl). Pockets were cast into the membrane as described in Ref.^40^. Embryos were then mounted into the pockets for imaging.

#### Immunofluorescence staining

Embryos were washed twice with PBS, fixed using 1% PFA and 0.1% Triton X-100 (Sigma, T8787) in PBS for 30 min at room temperature, and then washed again three times with PBS. All these steps were carried out in IBIDI slides coated with 0.5% Agar. After washing, the embryos were transferred to uncoated IBIDI slides containing blocking buffer (PBS added with 0.1% Triton X-100, 3% BSA (Sigma, 9647) and 5% Normal Goat Serum) and incubated over night at 4°C. After blocking, the embryos were incubated over night at 4°C with primary antibodies diluted in blocking buffer. Afterwards, the embryos were washed once with 0.1% BSA and 0.1% Triton X-100 in PBS, incubated for 1h at room temperature with secondary antibodies and DAPI (Invitrogen, D3751; 1:2000) diluted in blocking buffer, and washed again with 0.1% BSA and 0.1% Triton X-100 in PBS. Finally, the embryos were mounted individually in 0.5 ul drops of 0.1% BSA in PBS and placed in LabTek chambers covered with oil. Primary antibodies against CDX2 (Biogenex, NC9471689; 1:150), SOX2 (Sigma, SAB3500187; 1:50) were used in this study. For the secondary staining, antibodies targeting mouse immunoglobulin coupled to Alexa Fluor 488 (Life Technologies, A21202), Alexa Fluor 555 (Life Technologies, A31570), rabbit immunoglobulin-coupled Alexa Fluor 546 (Invitrogen, A10040) were used. Finally, Alexa Fluor 633-coupled Phalloidin (Invitrogen, R415; 1:50) was used to stain filamentous Actin.

## Code availability

The GPU-accelerated spectral analysis code can be accessed at https://github.com/prevedel-lab/brillouin-gpu-acceleration.

## Data availability

The raw datasets generated and/or analysed during the current study will be made publicly available at the point of publication.

## REFERENCES

1. Engler, A. J., Sen, S., Sweeney, H. L. & Discher, D. E. Matrix Elasticity Directs Stem Cell Lineage Specification. Cell 126, 677–689 (2006).

2. Krieg, M. et al. Atomic force microscopy-based mechanobiology. Nat. Rev. Phys. (2018) doi:10.1038/s42254-018-0001-7.

3. Hochmuth, R. M. Micropipette aspiration of living cells. J. Biomech. 33, 15–22 (2000).

4. Kennedy, B. F., Wijesinghe, P. & Sampson, D. D. The emergence of optical elastography in biomedicine. Nat. Photonics 11, 215 (2017).

5. Palombo, F. & Fioretto, D. Brillouin Light Scattering: Applications in Biomedical Sciences. Chem. Rev. 119, 7833–7847 (2019).

6. Prevedel, R., Diz-Muñoz, A., Ruocco, G. & Antonacci, G. Brillouin microscopy: an emerging tool for mechanobiology. Nat. Methods 16, 969–977 (2019).

7. Antonacci, G. et al. Recent progress and current opinions in Brillouin microscopy for life science applications. Biophys. Rev. 12, 615–624 (2020).

8. Scarcelli, G. & Yun, S. H. Confocal Brillouin microscopy for three-dimensional mechanical imaging. Nat Phot. 2, 39–43 (2008).

9. Scarcelli, G. et al. Noncontact three-dimensional mapping of intracellular hydromechanical properties by Brillouin microscopy. Nat. Methods 12, 1132–1134 (2015).

10. Mattana, S., Caponi, S., Tamagnini, F., Fioretto, D. & Palombo, F. Viscoelasticity of amyloid plaques in transgenic mouse brain studied by Brillouin microspectroscopy and correlative Raman analysis. J. Innov. Opt. Health Sci. 10, 1742001 (2017).

11. Chan, C. J., Bevilacqua, C. & Prevedel, R. Mechanical mapping of mammalian follicle development using Brillouin microscopy. Commun. Biol. 4, 1133 (2021).

12. Scarcelli, G. & Yun, S. H. In vivo Brillouin optical microscopy of the human eye. Opt. Express 20, 9197–9202 (2012).

13. Schlüßler, R. et al. Mechanical Mapping of Spinal Cord Growth and Repair in Living Zebrafish Larvae by Brillouin Imaging. Biophys. J. 115, 911–923 (2018).

14. Bevilacqua, C., Sánchez-Iranzo, H., Richter, D., Diz-Muñoz, A. & Prevedel, R. Imaging mechanical properties of sub-micron ECM in live zebrafish using Brillouin microscopy. Biomed. Opt. Express 10, 1420 (2019).

15. Palombo, F., Madami, M., Stone, N. & Fioretto, D. Mechanical mapping with chemical specificity by confocal Brillouin and Raman microscopy. Analyst 139, 729–733 (2014).

16. Koski, K. J., Akhenblit, P., McKiernan, K. & Yarger, J. L. Non-invasive determination of the complete elastic moduli of spider silks. Nat. Mater. 12, 262 (2013).

17. Yun, S. H. & Chernyak, D. Brillouin microscopy: Assessing ocular tissue biomechanics. Curr. Opin. Ophthalmol. 29, 299–305 (2018).

18. Remer, I., Shaashoua, R., Shemesh, N., Ben-Zvi, A. & Bilenca, A. High-sensitivity and high-specificity biomechanical imaging by stimulated Brillouin scattering microscopy. Nat. Methods 0–18 (2020) doi:10.1038/s41592-020-0882-0.

19. Ballmann, C. W. et al. Stimulated Brillouin Scattering Microscopic Imaging. Sci. Rep. 5, 18139 (2016).

20. Zhang, J., Fiore, A., Yun, S.-H., Kim, H. & Scarcelli, G. Line-scanning Brillouin microscopy for rapid non-invasive mechanical imaging. Sci. Rep. 6, 35398 (2016).

21. Strnad, P. et al. Inverted light-sheet microscope for imaging mouse pre-implantation development. Nat. Methods 13, 139–142 (2015).

22. Elsayad, K. et al. Mapping the subcellular mechanical properties of live cells in tissues with fluorescence emission–Brillouin imaging. Sci. Signal. 9, rs5–rs5 (2016).

23. Meng, Z., Traverso, A. J. & Yakovlev, V. V. Background clean-up in Brillouin microspectroscopy of scattering medium. Opt. Express 22, 5410–5415 (2014).

24. Caponi, S., Fioretto, D. & Mattarelli, M. On the actual spatial resolution of Brillouin Imaging. Opt. Lett. 45, 1063 (2020).

25. Antonacci, G., Foreman, M. R., Paterson, C. & Török, P. Spectral broadening in Brillouin imaging. Appl. Phys. Lett. 103, 5–8 (2013).

26. Nikolić, M. & Scarcelli, G. Long-term Brillouin imaging of live cells with reduced absorption-mediated damage at 660nm wavelength. Biomed. Opt. Express 10, 1567 (2019).

27. Martin, A. C. The Physical Mechanisms of Drosophila Gastrulation: Mesoderm and Endoderm Invagination. Genetics 214, 543–560 (2020).

28. Bhide, S. et al. Mechanical competition alters the cellular interpretation of an endogenous genetic program. J. Cell Biol. 220, (2021).

29. Nishikata, T., Hibino, T. & Nishida, H. The Centrosome-Attracting Body, Microtubule System, and Posterior Egg Cytoplasm Are Involved in Positioning of Cleavage Planes in the Ascidian Embryo. Dev. Biol. 209, 72–85 (1999).

30. Liu, T. L. et al. Observing the cell in its native state: Imaging subcellular dynamics in multicellular organisms. Science (80-.). 360, (2018).

31. Krzic, U., Gunther, S., Saunders, T. E., Streichan, S. J. & Hufnagel, L. Multiview light-sheet microscope for rapid in toto imaging. Nat. Methods 9, 730–733 (2012).

32. Vergara, H. M. et al. Whole-body integration of gene expression and single-cell morphology. Cell 2020.02.26.961037 (2021) doi:10.1016/j.cell.2021.07.017.

## METHODS ONLY REFERENCES

33. Bakhshandeh, S. et al. Optical quantification of intracellular mass density and cell mechanics in 3D mechanical confinement. Soft Matter 17, 853–862 (2021).

34. Przybylski, A., Thiel, B., Keller-Findeisen, J., Stock, B. & Bates, M. Gpufit: An open-source toolkit for GPU-accelerated curve fitting. Sci. Rep. 7, 15722 (2017).

35. Bulgakova, N. A., Grigoriev, I., Yap, A. S., Akhmanova, A. & Brown, N. H. Dynamic microtubules produce an asymmetric E-cadherin-Bazooka complex to maintain segment boundaries. J. Cell Biol. 201, 887–901 (2013).

36. Campos-Ortega, J. A. & Hartenstein, V. The Embryonic Development of Drosophila melanogaster. (Springer Berlin Heidelberg, 1997). doi:10.1007/978-3-662-22489-2.

37. Gomez, J. M., Chumakova, L., Bulgakova, N. A. & Brown, N. H. Microtubule organization is determined by the shape of epithelial cells. Nat. Commun. 7, 13172 (2016).

38. Kölsch, V., Seher, T., Fernandez-Ballester, G. J., Serrano, L. & Leptin, M. Control of Drosophila gastrulation by apical localization of adherens junctions and RhoGEF2. Science 315, 384–6 (2007).

39. Robin, F. B. et al. Time-Lapse Imaging of Live Phallusia Embryos for Creating 3D Digital Replicas: Movie 1. Cold Spring Harb. Protoc. 2011, pdb.prot065847 (2011).

40. Reichmann, J., Eguren, M., Lin, Y., Schneider, I. & Ellenberg, J. Live imaging of cell division in preimplantation mouse embryos using inverted light-sheet microscopy. in 279–292 (2018). doi:10.1016/bs.mcb.2018.03.030.

41. Chan, C. J. et al. Hydraulic control of mammalian embryo size and cell fate. Nature 571, 112–116 (2019).

