## Supplementary Information for "High-resolution line-scan Brillouin microscopy for live-imaging of mechanical properties during embryo development"

### SUPPLEMENTARY FIGURES

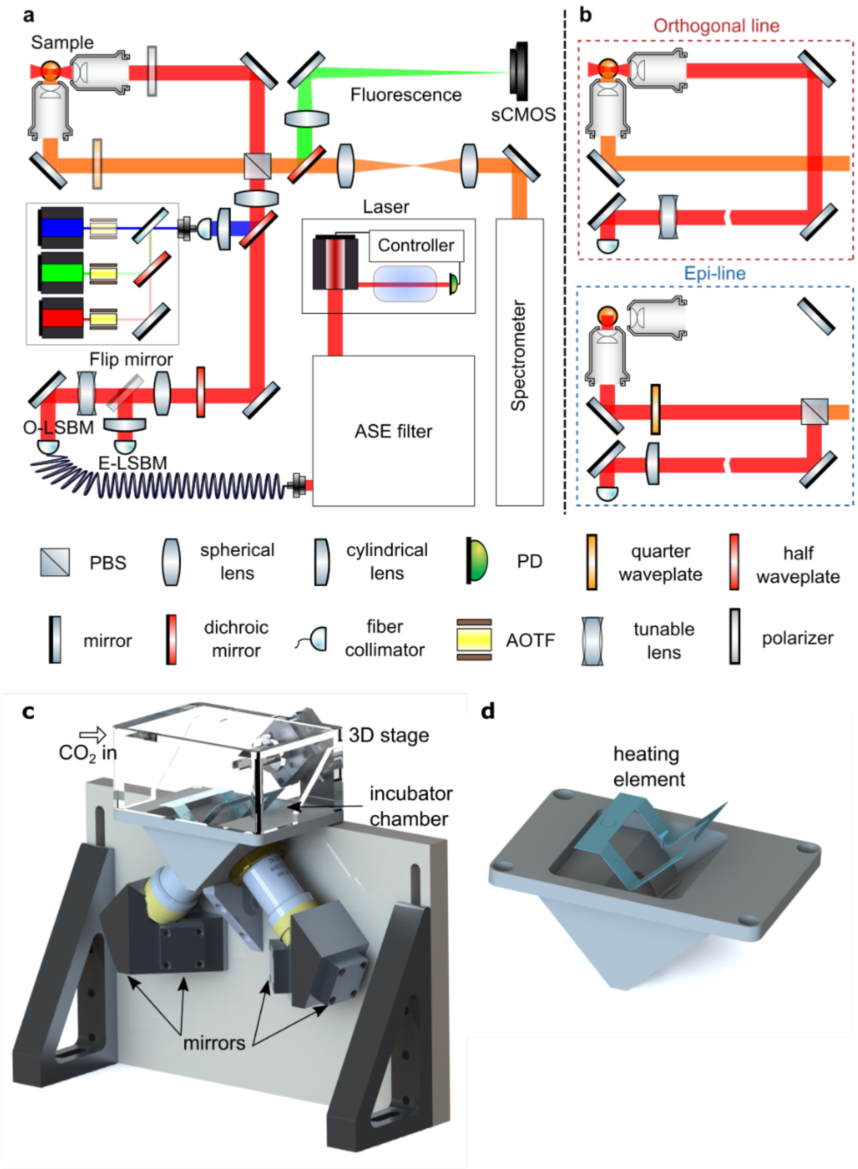


**SI Figure 1: Optical and mechanical design of the LSBM**
**(a)** Conceptual schematic of the optical layout including both the fluorescence and Brillouin imaging modalities. The spectrometer and ASE filter are further detailed in **SI Figs. 2** and **3**, respectively. **(b)** Optical path for the Brillouin illumination and detection in the orthogonal (top) and epi line (bottom) configuration. Note that switching between these configurations can be done by flipping the mirror after the tunable lens, adding/removing the PBS (mounted on a magnetic base to ease this operation) and rotating the half and quarter waveplates appropriately. **(c)** 3D rendering of the mechanical design. The objectives are arranged in an inverted 90 deg V-configuration; light is delivered (and collected) to the objectives by two mirrors that preserve image orientation. **(d)** Enlarged view of the immersion chamber, showing the heating element inset used for temperature control. See **Methods** for further details.

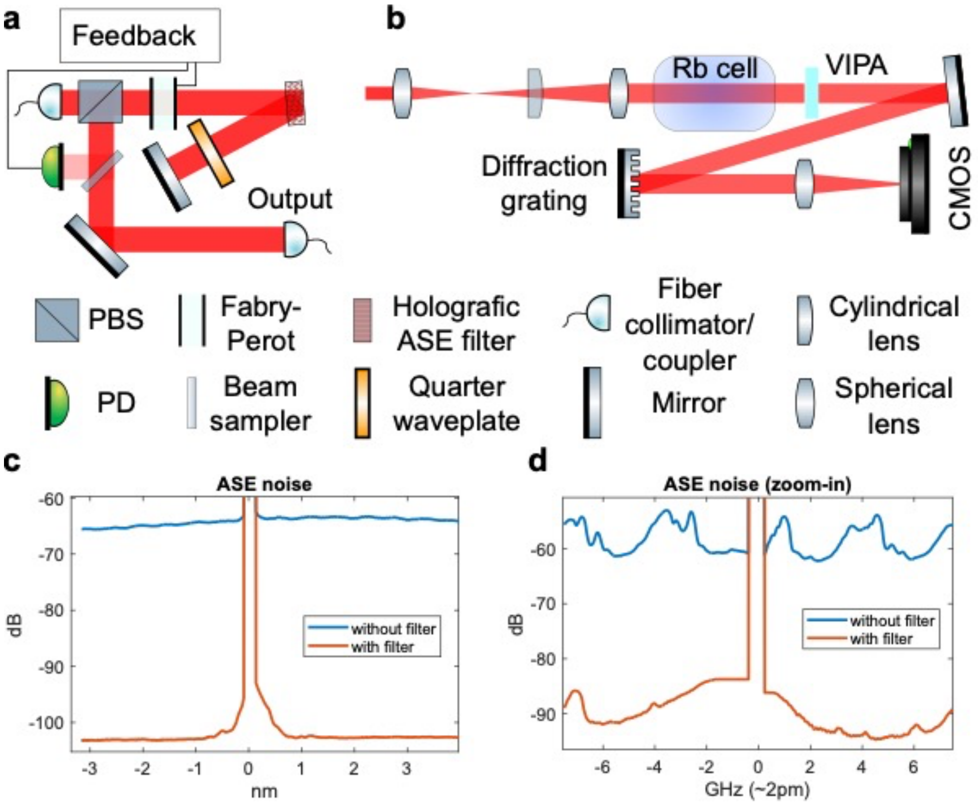


**SI Figure 2: Characterisation and filtering of Amplified Spontaneous Emission (ASE)** **(a)** Optical layout of the double-pass ASE filter employed. To stabilize the Fabry-Perot (FP) based filter over a long time a feedback-loop is realized via a photodiode, custom-written Arduino software and piezo that adjusts the FP mirror spacing. **(b)** Optical setup used to measure the ASE **(c,d)** Plots of ASE spectrum with and without filter few nm (c) and few GHz (d) away from the laser line. The filter suppresses the total ASE (integrated over the whole measured range) by more than 40dB. See **Methods** for further details.

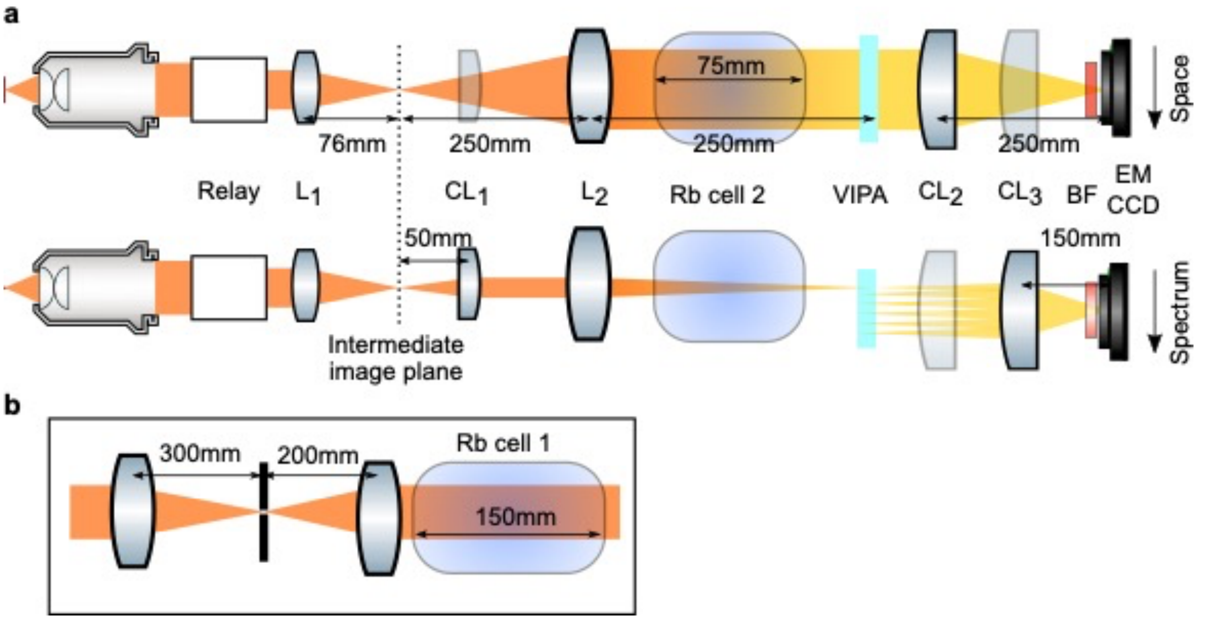


**SI Figure 3: Optical design of the BLSM spectrometer**
**(a)** Orthogonal views of the optical components in the BLSM spectrometer with relevant distances; top shows the plane determined by the illumination line and the optical axis of the detection objective, where the optics relay to the VIPA, and appropriately magnify, the BFA of the objective; bottom shows the plane perpendicular to the illumination line, where the optics relay to the VIPA, and appropriately magnify, the image plane. **(b)** Details of the relay shown in panel **a**.

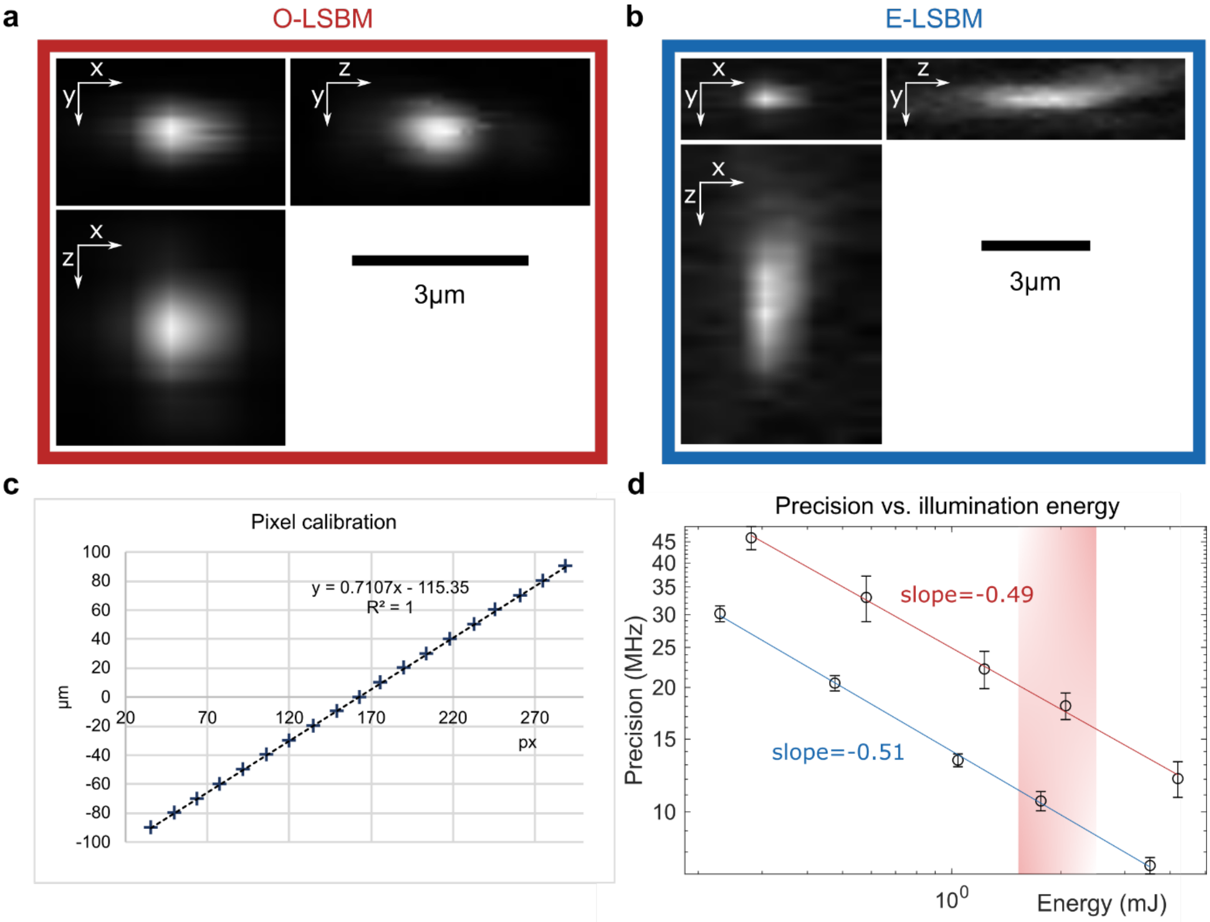


**SI Figure 4: Optical characterisation and precision measurement of the BLSM spectrometer.**
Orthogonal views of a 0.5um bead (Tetraspeck, Thermo Fischer) to characterise the optical resolution in the orthogonal **(a)** and epi line **(b)** geometries. **(c)** Spatial calibration (µm to pixel) acquired by translating a bead, with steps of 10 µm, along the x direction and determining its position on the camera. **(d)** Precision vs. illumination energy for the orthogonal (red) and epi (blue) line geometries. The fit shows a square root dependence, as expected in shot noise limited conditions. The shaded red region indicates the typical imaging conditions used in the experiments. See **Methods** for details.

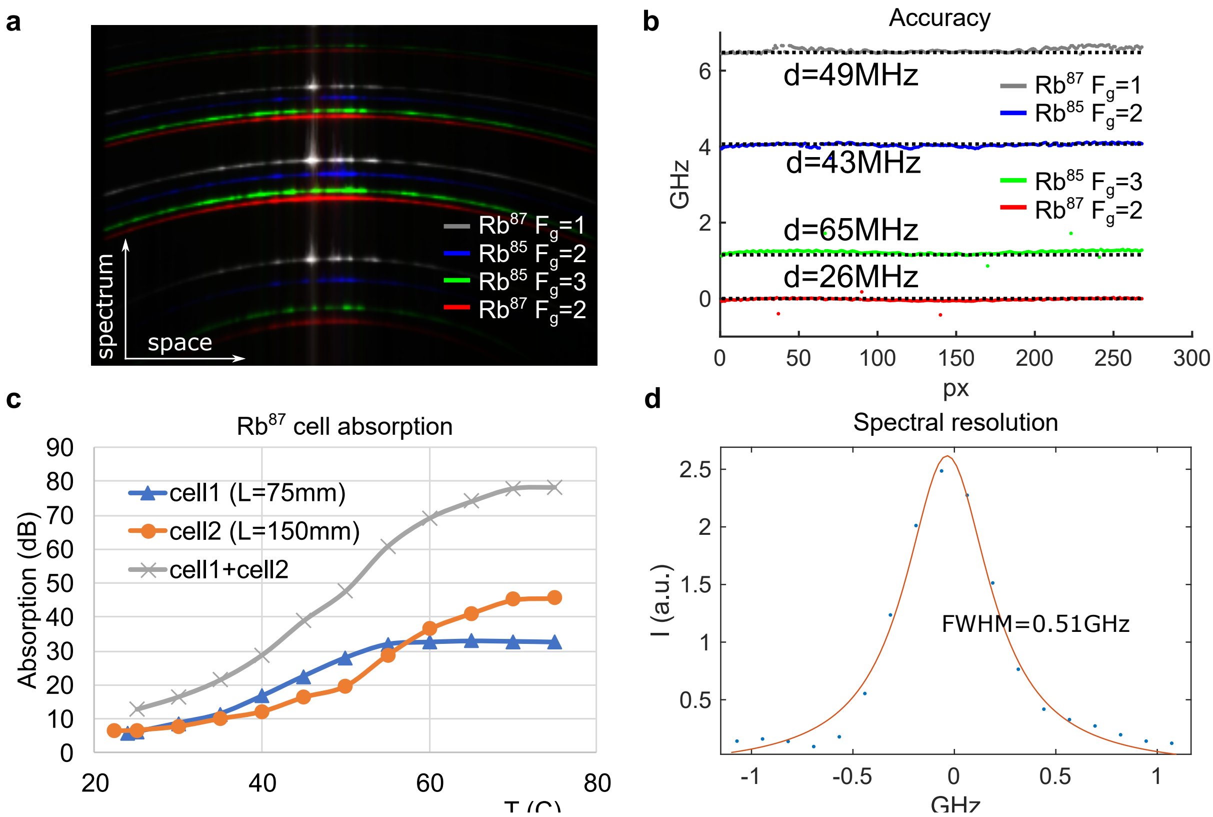


**SI Figure 5: Spectral characterisation of the BLSM spectrometer.**

**(a)** Acquired, raw spectra of the 4 hyperfine D2 spectral lines of $Rb^{85}$ and $Rb^{87}$ as measured by the LSBM spectrometer. **(b)** Frequency shift of the Rb lines in panel **a** after applying the reconstruction pipeline (dotted lines represent the exact values measured with a frequency counter); d is the $L^1$ distance between the measured and the exact values divided by the number of points i.e. the average discrepancy from the true frequency; the FSR is measured to be 15.15GHz by minimizing the sum of d for the 4 Rb lines. **(c)** Suppression of Rayleigh-scattered light from a 150mm long (orange line) and 75mm long (blue line) Rb cell as a function of the temperature. **(d)** Spectral resolution of the spectrometer, as measured by the FWHM of the laser line, after applying the reconstruction pipeline.

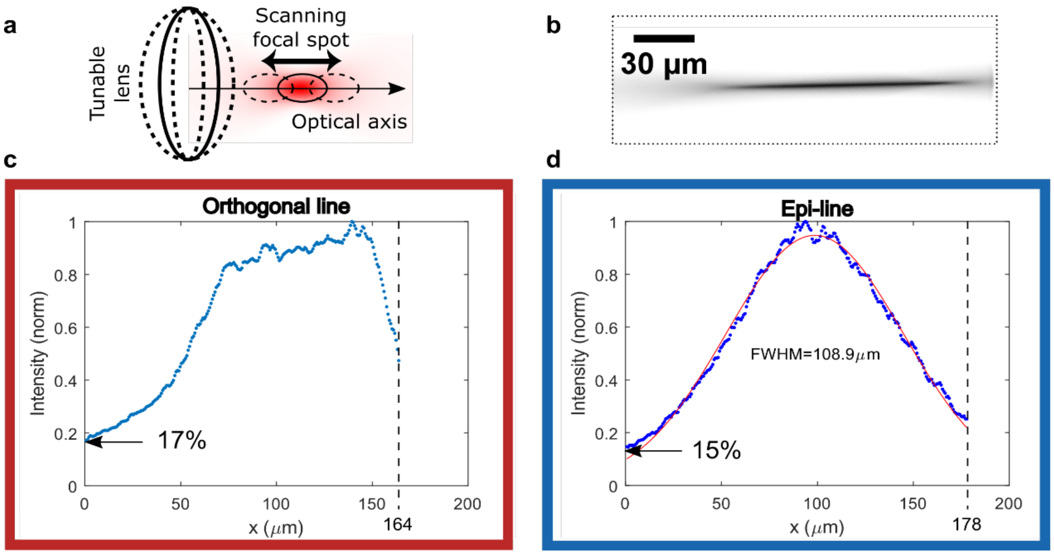


**SI Figure 6: Characterisation of the Brillouin illumination profile for both orthogonal and epi line geometries.**

**(a)** Illustration of axial scanning by means of a tunable lens in order to extend the illumination depth and thus FOV in the orthogonal geometry. The curvature of the tunable lens can be controlled with an external voltage, thus shifting the focus spot in the sample. **(b)** Fluorescence image of the extended illumination line in the orthogonal geometry, acquired by imaging the fluorescence signal from ICG dye with a camera in the intermediate image plane (dashed line in **SI Fig. 3**). **(c,d)** Signal intensity profile along the FOV (measured as the intensity of the Stokes/anti-Stokes spectral peaks in water) for the **(c)** epi line and **(d)** orthogonal line geometries, respectively. Note that this is a proxy for the illumination profile under the assumption that there are no spatially dependent losses in the spectrometer. Also note that a relative intensity value of >15% is generally sufficient for reliable Brillouin peak fitting/detection.

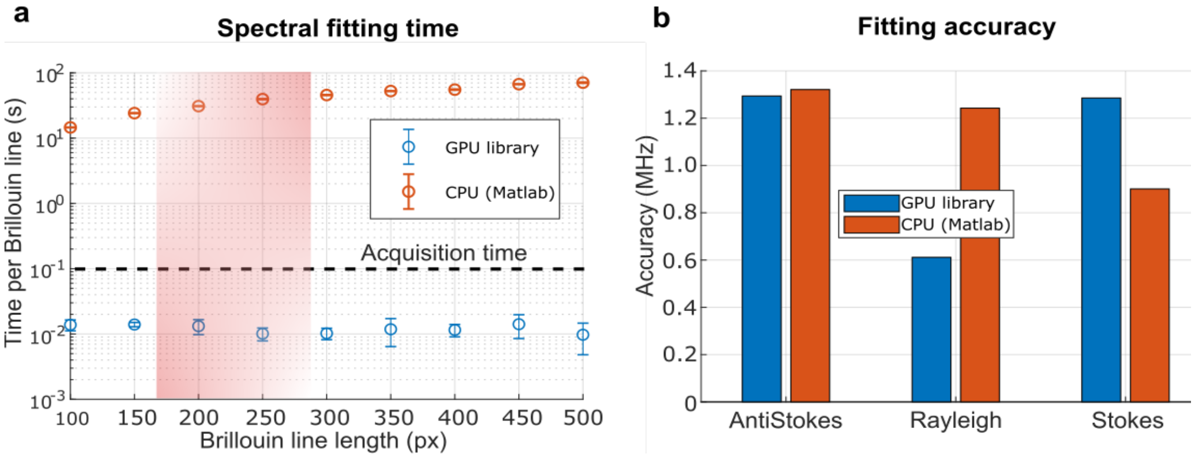


**SI Figure 7: Performance of the GPU-accelerated spectral analysis pipeline.**

**(a)** Brillouin spectrum fitting time as a function of image size (equal to number of independent spectra), comparing standard, CPU-based fitting routines (Matlab) and our custom GPU-accelerated library. Total analysis time is generally >1000-fold improved with GPU acceleration, making the processing time smaller than the acquisition time. The shaded red region shows the typical length of the Brillouin line in pixels. Note most error bars are smaller than data points. **(b)** Fitting accuracy between the CPU- and GPU-based pipelines, showing comparable performance. Note that the accuracy uncertainty (~1MHz) is negligible compared to the spectral fitting precision based on realistic, noisy data (~10-20MHz).

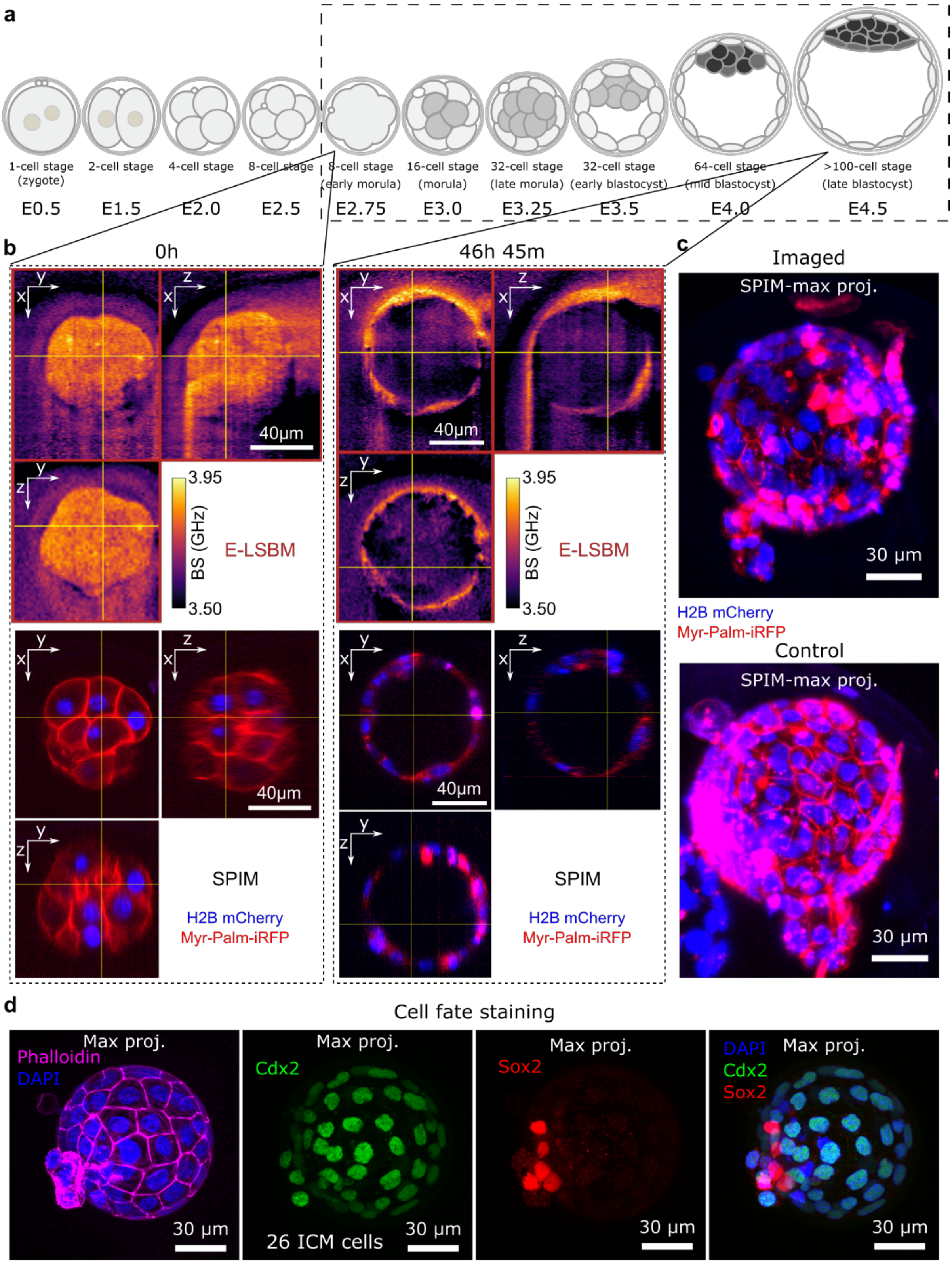


**SI Figure 8: LSBM allows Brillouin time-lapse imaging of developing mouse embryos over ~2 days.**

**(a)** Timeline of mouse embryo development from the one cell stage to late blastocyst. The dashed rectangle encloses the developmental window that was imaged. **(b)** Exemplary orthogonal views of Brillouin volumes (top, acquired in the O-LSBM) and SPIM volumes (bottom) of a single mouse embryo at the beginning (left) and at end of the timelapse (right, 46h45m after the first timepoint). The acquisition time for one volume in the Brillouin modality was ~11-17min, the time interval between volumes is between 77 to 92 minutes. Representative result from n=4 embryos. **(c)** Top: SPIM volume (maximum intensity projection -MIP) of an embryo that underwent Brillouin time-lapse imaging, taken at the end of the timelapse. Bottom: SPIM image (MIP) of a control embryo (taken at the same time as the embryo in top panel), that was in the same imaging chamber but not imaged by Brillouin or SPIM, showing qualitatively similar morphology. **(d)** To further confirm embryo viability, we fixed the embryos after imaging and stained them for cell fate. MIPs through the volume where the outer trophectoderm cells are CDX2-positive (green), a marker for trophectoderm cell fate. The inner cell mass are SOX2-positive (red) and CDX2-negative, indicating proper epiblast fate in the ICM at late blastocyst stage. The number of cells in the ICM is 26, consistent with the reported values in literature [1].

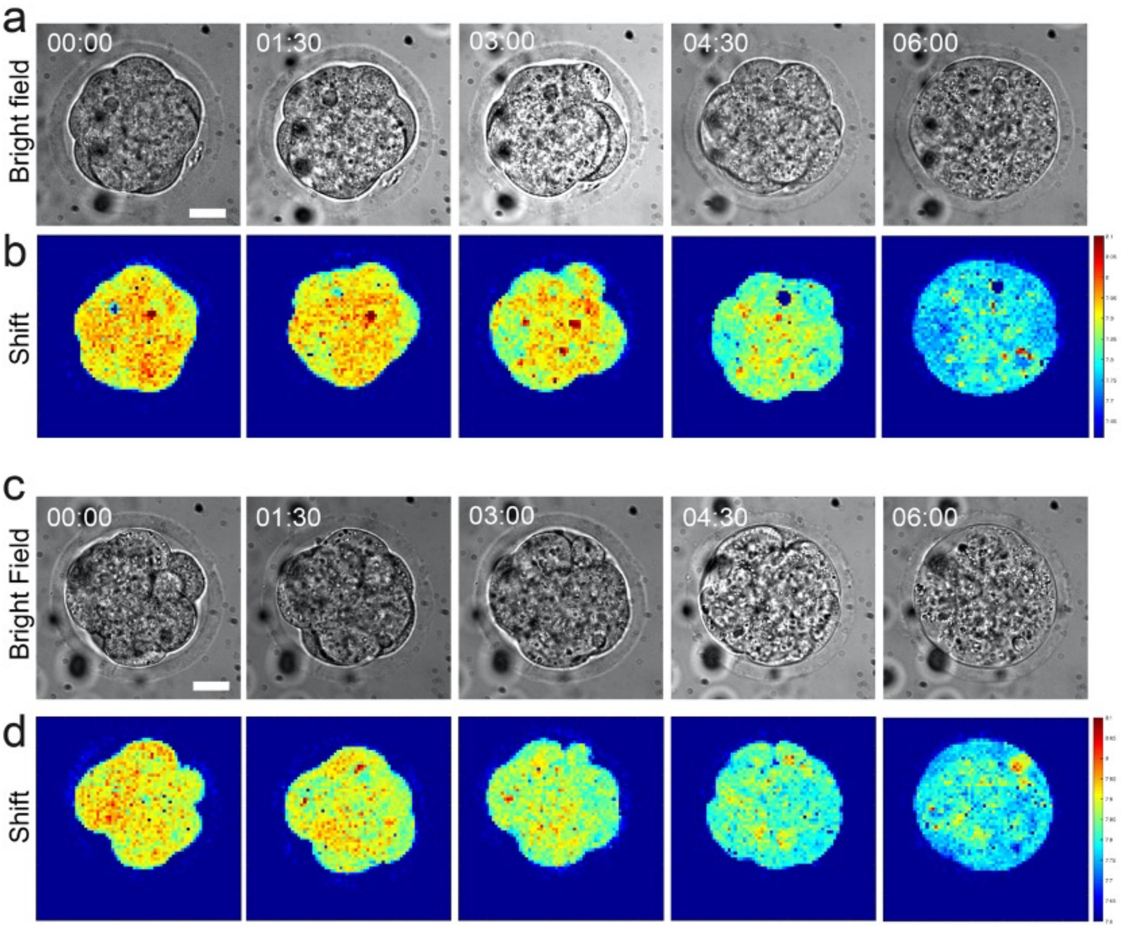


**SI Figure 9: Embryos are not viable under 532-nm confocal Brillouin imaging. a.** Bright-field images of an 8-cell stage mouse embryo that eventually undergoes cell death. **b.** Images of Brillouin shift for the same embryo in (**a**), showing a loss of Brillouin signals within 6 hours, which is a clear sign of embryo death. The embryo also appears round and decompacted, a sign of abnormality. **c.** Another representative embryo undergoing cell death under the same imaging condition. Time is shown in h:mm. Scale bar = 20 µm.

### SUPPLEMENTARY REFERENCES

1. C. J. Chan, M. Costanzo, T. Ruiz-Herrero, G. Mönke, R. J. Petrie, M. Bergert, A. Diz-Muñoz, L. Mahadevan, and T. Hiiragi, "Hydraulic control of mammalian embryo size and cell fate," Nature **571**, 112–116 (2019).

### SUPPLEMENTARY VIDEOS

**Supplementary Video 1: Brillouin shift map and fluorescence time-lapse from a *Drosophila* embryo undergoing ventral furrow formation (VFF).** Time-lapse of a transgenic *Drosophila* embryo expressing Gap43-mCherry to fluorescently label membranes, acquired from the onset of ventral furrow formation until invagination of the mesoderm is completed. Both fluorescent and Brillouin channels were re-sliced (left, bilinear interpolation), and the resulting images were Median-projected (two slices). The initiation of apical constriction was set as time-point '0'. The total volume of ~22µm x 170µm x 71µm was acquired in ~2min with a pixel size of 0.7µm x 1.5µm x 1.5µm. Note that the acquisition of the data shown in the video required <20s per timepoint. Time is in minutes, scale-bar is 30 µm.

**Supplementary Video 2: Brillouin shift map and fluorescence time-lapse from a *Drosophila* embryo undergoing posterior midgut invagination (PMG).** Time-lapse of a transgenic *Drosophila* embryo expressing Gap43-mCherry to fluorescently label membranes, acquired from the onset of posterior midgut invagination until invagination of both endoderm and pole cells is completed. Both fluorescent and Brillouin channels were Median-projected (two slices). The initiation of apical constriction was set as time-point '0'. The total volume of ~83µm x 141µm x 46µm was acquired in ~2min with a pixel size of 0.7µm x 1.5µm x 2.5µm. Note that the acquisition of the data shown in the video required <15s per timepoint. Time is in minutes, scale-bar is 30 µm.

**Supplementary Video 3: Single plane from the Brillouin shift and fluorescence map time-lapse (~14h) of a late tailbud I *Phallusia mammillata* embryo showing vacuoles formation and fusion.** Time-lapse (~14h) of a late tailbud I *Phallusia mammillata* embryo showing both fluorescence channel (left), where membranes are stained with FM4-64 dye, and Brillouin shift map (right). Note the vacuoles forming in the tail are clearly distinguishable by their lower Brillouin shift. The total volume of ~190µm x 190µm x 110µm with a pixel size of 0.7µm x 2µm x 3µm was acquired in ~9min.

**Supplementary Video 4: Maximum intensity projection from the Brillouin shift and fluorescence map time-lapse (~14h) of a late tailbud I *Phallusia mammillata* embryo showing vacuoles formation.** Time-lapse (~14h) of a late tailbud I *Phallusia mammillata* embryo showing both fluorescence channel (left), where membranes are stained with FM4-64 dye, and Brillouin shift map (right). Note the visible tail unfolding which is a characteristic of the late tailbud II stage, and thus an indicator for successful development. The total volume of ~190µm x 190µm x 110µm with a pixel size of 0.7µm x 2µm x 3µm was acquired in ~9min.
